# Primary culture of high grade serous ovarian cancer cells, selection and derivation of three cell lines

**DOI:** 10.1101/2024.08.15.607946

**Authors:** Maria Vias, Carolin M Sauer, Ian Goldlust, Chandra Chilamakuri, Douglas Hall, Teodora Goranova, Elke Van Oudenhove, Gabriel Funingana, Anna Piskorz, Elizabeth Moore, Helena M. Earl, James D. Brenton

## Abstract

Cell line models for high grade serous ovarian cancer (HGSOC) are limited in number and are poorly clinically annotated. Consequently, existing models often fail to recapitulate common features of HGSOC, inhibiting mechanistic and therapeutic discovery. We generated and characterised three spontaneously immortalized continuous HGSOC cell lines named CIOV1, CIOV2, and CIOV3 and confirmed that each cell line retained the genomic and pathologic characteristics of its parental tumour. We show that subclonal cell populations present at initiation, expanded and contracted during the culturing process before converging on a stable immortalized line. These lines are new valuable models to study acquired chemoresistance of HGSOC.

## Introduction

Although high grade serous ovarian cancer (HGSOC) is the most frequent histological ovarian cancer subtype, long-term cellular models of this disease are very limited but urgently needed. HGSOC is molecularly characterized by ubiquitous *TP53* mutation (Ahmed et al., 2010), profound genetic heterogeneity and copy number aberrations leading to specific copy number signatures (Macintyre et al., 2018; Drews et al., 2022). Approximately 50% of cases are predicted to show homologous recombination deficiency (HRD) and between 10% to 15% of patients have germline mutations in *BRCA1* or *BRCA2*.

We and others have recently developed long-term HGSOC organoid models (Kopper et al., 2019; Hoffmann et al., 2020; Senkowski et al., 2023; Vias et al., 2023) but there are still significantly fewer continuous cell lines, as defined as able to grow over 100 population doublings (Geraghty et al., 2014), than those from other tumour types as reflected in publicly available cell banks (ECACC and ATCC). Furthermore, only a fraction of purported HGSOC cell lines are correctly characterized and accuracy is inversely associated with frequency of use (Domcke et al., 2013). As a result, new cell lines have been generated (Fleury et al., 2015; Ince et al., 2015; Kreuzinger et al., 2015; Thu et al., 2017). However, many important variables remain unresolved particularly key questions about success rate, time needed in culture or how evolution and selection may alter the final derived continuous line. These data are critical to inform model selection, use and future derivation of HGSOC lines.

The primary aim of this work was to systematically determine cell survival of HGSOC tumours following typical culture methods based on established guidelines (Geraghty et al., 2014) and to determine whether selection significantly alters genomic representation of the parental tumour.

## Materials and methods

### Patient data

Clinical data and tissue samples for 93 patients were collected on the prospective Cambridge Translational Cancer Research Ovarian Study 04 (CTCR-OV04) with Research Ethics Committee approval 08/H0306/61. Patients provided written, informed consent for participation in this study and for the use of their donated tissue for the laboratory studies carried out in this work.

### Sample collection and processing

#### Solid tumour collection and processing

During surgery the tissues were placed in DMEM/F12 (Gibco) medium supplemented with 10 mmol/l Hepes, 5% FBS and 50 µg/ml gentamicin and kept on ice. Tissues were received within 1 h of surgical excision. Solid tumour ovarian cancer samples were minced mechanically into 2 mm to 5 mm pieces using scalpels and added to dissociation medium containing DMEM/F12 supplemented with 50 μg/ml gentamicin, 1.5% Bovine Serum Albumin Fraction V, 5 μg/mL insulin, 1 mg/mL collagenase A and 100 U/ml hyaluronidase. Tissues were incubated for 16 h in a rotatory shaker at 80 rpm and 37 °C. After the incubation, cells were spun down and washed with DMEM/F12 medium. The cells were further disaggregated with 0.25% trypsin-EDTA, 5 mg/ml dispase and 0.1 mg/ml DNase (Sigma) treatment for 4 min. Erythrocytes were lysed using ammonium chloride solution (StemCell Technologies). Viable cells were counted with Trypan Blue exclusion dye.

#### Ascitic/pleural fluid collection and processing

Ascites fluid was drained from patients by inserting a routine peritoneal drainage catheter if this was a therapeutic procedure or a needle for diagnostic purposes. All procedures were done routinely as part of standard patient care. The fluid was collected by gravity into sterile Duran bottles and centrifuged into plastic conical tubes at 450×*g* for 10 min. Cells were then washed with DMEM/F12 medium and centrifuged at 400×*g* for 5 min.

### Culture conditions

All cell line work was carried out following published guidelines for the use of cell lines in biomedical research (Geraghty et al., 2014). Single cell suspensions were grown in DMEM/F12 medium supplemented with 10% FBS at 37 °C and 5% CO_2_ using standard cell culture dishes. In all cases, primary cells were cultured in antibiotic-free conditions and were routinely tested against mycoplasma infection. Cells were monitored twice a week for morphological changes using an inverted microscope and cultures were stopped when cells were observed to be dead or senescent. Cells were passaged when 80% confluency was reached and considered to be continuous cell lines when 100 population doublings were achieved.

### Survival analyses

Survival distributions were compared using the logrank test using the R statistical language. Cultures were considered not viable when the flasks contained only dead floating cells or were overtaken by fibroblast-like (Supplementary Figure A left panel) or senescent cells (Supplementary Figure A right panel). For the analysis cell lines were censored when frozen down.

### Short Tandem Repeat (STR) profiling

DNA extraction was performed using the DNeasy Blood & Tissue Kit (QIAGEN) according to manufacturer’s instructions. STR genotyping was performed using the GenePrint® 10 kit (Promega).

### Immunohistochemistry

Haematoxylin and Eosin (H&E) slides were stained according to the Harris H&E staining protocol and using a Leica ST5020 multi-stainer instrument. Staining for p53 and vimentin was done using the Leica Bond Max fully automated IHC system. Briefly the slides were retrieved using ER1 (Leica Microsystems) for 30 min, and p53 monoclonal antibody (clone DO-7) was employed at a 1/50 dilution for 30 min. Bond™ Polymer Refine Detection System (Leica Microsystems) was used to visualise the brown precipitate from the chromogenic substrate, 3,3’-Diaminobenzidine tetrahydrochloride (DAB).

### Mutation analysis

#### Tagged amplicon sequencing (TAm-Seq)

The coding sequences of *TP53, PTEN, BRCA1, BRCA2, MLH1, MSH2, MSH6, PMS2, RAD51C, RAD51B, RAD51D*, and hot spots for *EGFR, KRAS, BRAF, PIK3CA* were sequenced using tagged amplicon sequencing on the Fluidigm Access Array 48.48 platform as previously described (Forshew et al., 2012). The libraries were sequenced on the MiSeq platform using paired-end 125bp reads. Variant calling from sequencing data was performed using an in-house analysis pipeline and IGV software as described previously (Thorvaldsdóttir et al., 2013). Synonymous and non-pathogenic nonsynonymous mutations or nonsynonymous mutations of unknown clinical significance were excluded.

#### Sanger sequencing

The *TP53* exons 2 to 11 were amplified as previously described (Sjöblom et al., 2006) with the following modifications: PCR reactions were performed in 25µl of water and universal primers M13 forward and reverse were incorporated into primer pairs. After amplification the samples were submitted to GATC Biotech AG for Sanger sequencing with M13 primers and mutational analysis was performed using Mutation Surveyor Software v4.0.4 (SoftGenetics) using default settings.

#### Digital PCR

Digital PCR was performed using the 12.765 digital chip from the Fluidigm Biomark microfluidic system according to the manufacturer’s instructions. Primers were designed spanning the *NF1* deletion (forward: TTTTGTTTACGAGCACAGATAACC and reverse: GAAACAGAAGATGACAGCAAAGAA). Digital PCR was also performed on the same samples using an assay for the p.R175H mutation in*TP53* as previously described (Schwarz et al., 2015). The proportion of tumour cells with the *NF1* deletion was calculated from the estimated counts for both the *NF1* deletion and mutant *TP53* (p.R175H).

### Shallow whole genome sequencing (sWGS)

Whole genome libraries were prepared using the TruSeq Nano Kit as previously described (Piskorz et al., 2016). Each library was quantified using the KAPA Library Quantification kit (kappa Biosystems) and 10 nM of each library was combined in a pool of 21 samples and sequenced on the Illumina HiSeq 4000 machine using single-end 150-bp reads. Reads were aligned against the human genome assembly GRCh37 using the BWA-MEM algorithm (v0.7.12). Duplicates were marked using the Picard Tool (v1.47) (“Picard toolkit,” n.d.) and copy number was assessed using Bioconductor package QDNAseq (v1.6.1) (Scheinin et al., 2014).

### Absolute copy number calling

Relative to absolute copy-number scaling was generated by Rascal (Sauer et al., 2021). Absolute copy-number and mutant allele frequency, identified using tagged amplicon, were combined in a probabilistic graphical modelling approach to infer absolute copy-number signatures as described previously (Macintyre et al., 2018).

### RNAseq and PROGENy analysis

Sequencing reads were mapped to the GRCh38 (hg38) reference genome using STAR aligner v2.7.6a. Quantified transcripts (gene-level) were imported into R using the tximport (Soneson et al., 2016) in preparation for downstream analyses. Low-abundance genes were filtered out, keeping only genes with at least 25 reads in at least two samples. Gene count normalisation and differential gene expression analyses were performed using DESeq2 v1.26.0 (Love et al., 2014). Genes with p-value < 0.05, FDR < 0.1 and a fold-change of at least ×1.5 were considered differentially expressed. PROGENy pathway activities were performed as previously described (Schubert et al., 2018; Holland et al., 2020) for each cell line using progeny R package v1.16.0 (“Pathway RespOnsive GENes for activity inference from gene expression,” n.d.).

### Cell proliferation and senescence

Before performing any assay, cells were counted using the Luna cell counter and viability was assessed using Trypan blue dye. For any given assay viability was always between 88-98%. Cells were seeded at 30,000 cells/well in 24-well plates and visualized using a live cell imaging system (IncuCyte™ ESSEN BioScience Inc, Ann Arbor, Michigan, USA). Cells were imaged every 3 h for several days to monitor cell confluency and morphological changes. Proliferation was plotted using the R statistical language. Senescence was assessed using the senescence cell staining kit (Sigma) according to manufacturer instructions.

### 3D cell growth

The ability of the cell lines to generate a three-dimensional structure was assessed using round-bottom ultra-low attachment (ULA) plates (Corning) and ECM support (Growth Factor Reduced BD Matrigel®). Cells (3,000 cells/well) were seeded in complete medium and incubated at 37 °C for 14 d before imaging.

### Wound-healing assay

The migration potential of the cell lines was assessed using the scratch assay method. Cells were grown to confluence in 96-well Essen Image Lock™ plates and wounded with the Wound Maker® device (IncuCyte™ ESSEN BioScience Inc.). Cells were monitored every 3 h for 24 hours.

### Chemosensitivity assays

Cells were seeded in 96-well plates (10,000 cells/well) and treated the next day with vehicle, cisplatin (0-320 µM) or paclitaxel (0-320 nM) for 96 h. Cells were washed with PBS, fixed with 3.3% TCA (w/v) for 1 h at 4 °C and stained with sulforhodamine B for 30 min. After washing cells were solubilised using 200 µl of Tris base solution (pH 10.5) for 30 min, and absorbance was measured at 510 nm using the spectrophotometer BMG PHERAstar FS. Drug-response data were fit with a four-parameter log-logistic function using the GR metrics package (Clark et al., 2017). GR value can be interpreted as the ratio between growth rates under treated and untreated conditions normalized to a single cell division.

### *In vivo* growth

Animal procedures were followed in compliance with the local AWERB, NACWO and UK Home Office regulations. Five million cells were re-suspended in 25 µl of DMEM/F12 media and mixed with 25 µl of Growth Factor Reduced BD Matrigel®. Cells were injected subcutaneously in both flanks of NSG female mice (two mice per cell line). Tumour formation was assessed weekly.

## Results

### *In vitro* survival of high grade serous ovarian cancer cells

A total of 114 primary tissue samples comprising 50 liquid specimens (ascites and pleural fluid) and 64 solid tumours were grown *in vitro*. We standardized culture conditions using same media (DMEM/F12 + 10% FBS) and plasticware. For some patients, more than one tissue type was collected (eg: omentum, ovary, ascitic fluid or different ascites time point collections). The success of cell culture experiments was ascertained using time to event and Kaplan Meier estimator analysis (Figure 1A). Of the 114 samples cultured, 13 primary cultures were excluded. Ten were not HGSOC in origin and three did not harbour a *TP53* mutation. From the remaining 101 HGSOC lines, 98 were uncensored and among these the median survival time was 22 days. 35% of the cultures failed within a few days of seeding (passage 0) and the remaining (65%) grew *in vitro* for 50 days (5-10 passages) becoming finite cell lines until the cells senesced (Supplementary Figure 1A). Although most cultures senesced before reaching passage 10, HGSOC finite cell lines were successfully generated for short-term experiments. Only three HGSOC lines that survived beyond passage 10 achieved continuous growth (Figure 1B).

**Figure 1.**
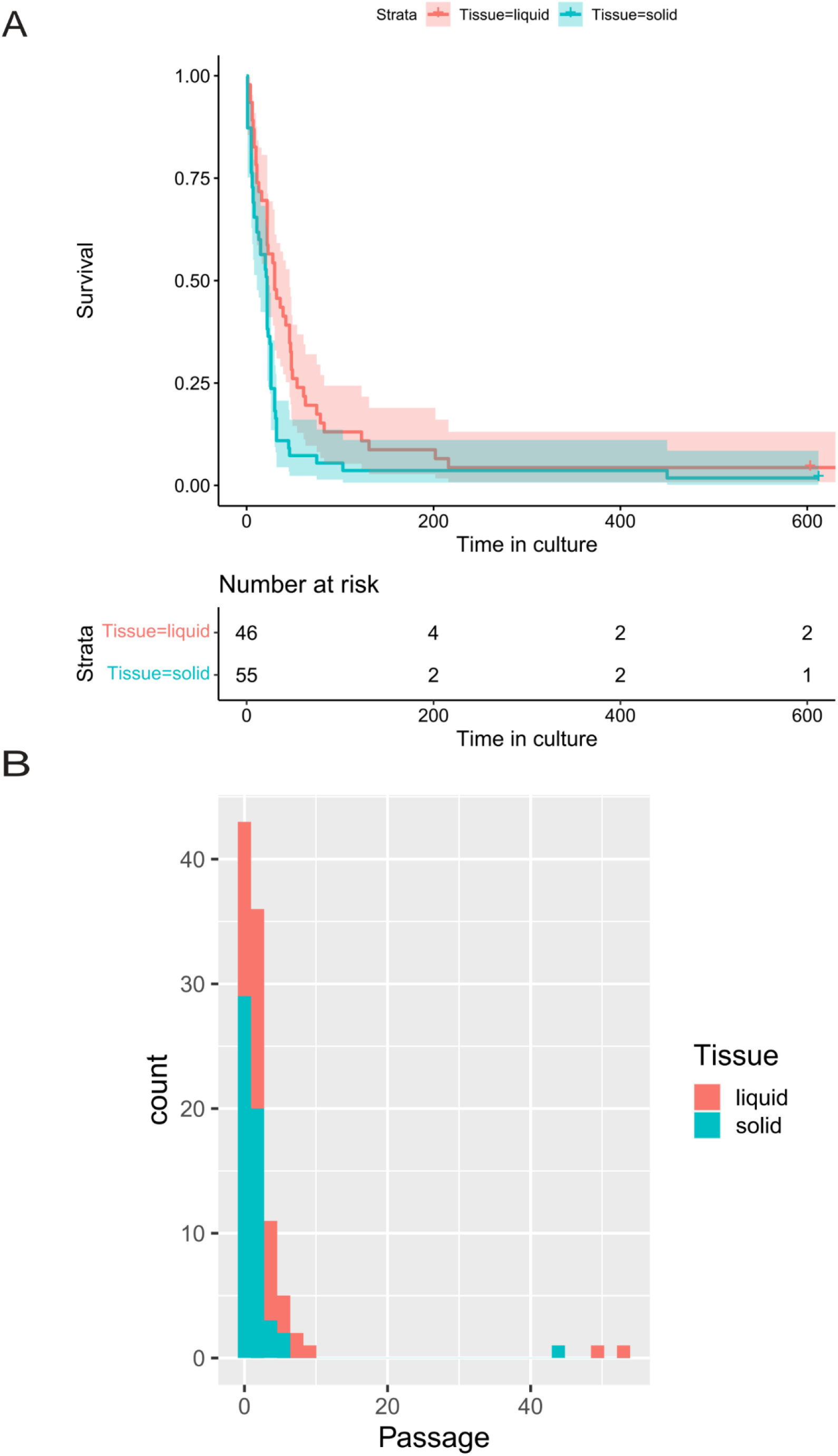
In vitro 2D survival of HGSOC cells. A. Kaplan Meier analysis of overall survival of the cells growing as 2D cultures. B. Passage number analysis of the cell lines growing as 2D cultures.

### Patient clinical history and cell line characterization

Our goal in generating these cell lines was to develop spontaneously immortalized, high-fidelity HGSOC *in vitro* models to facilitate research. To that end we describe here, in depth, the methods that we used to successfully establish three cell lines (CIOV1, CIOV2 and CIOV3) and provide definitive clinical, genomic, and phenotypic characterization to serve as a basis for future comparison.

#### Patient clinical data and culture of tumour cells

The patient’s treatment journey is summarised in Figure 2A, where tissue collection time is also shown. Patient OV04-97 was sensitive to chemotherapy, OV04-336 was intrinsically resistant and OV04-366 had acquired resistance.

**Figure 2.**
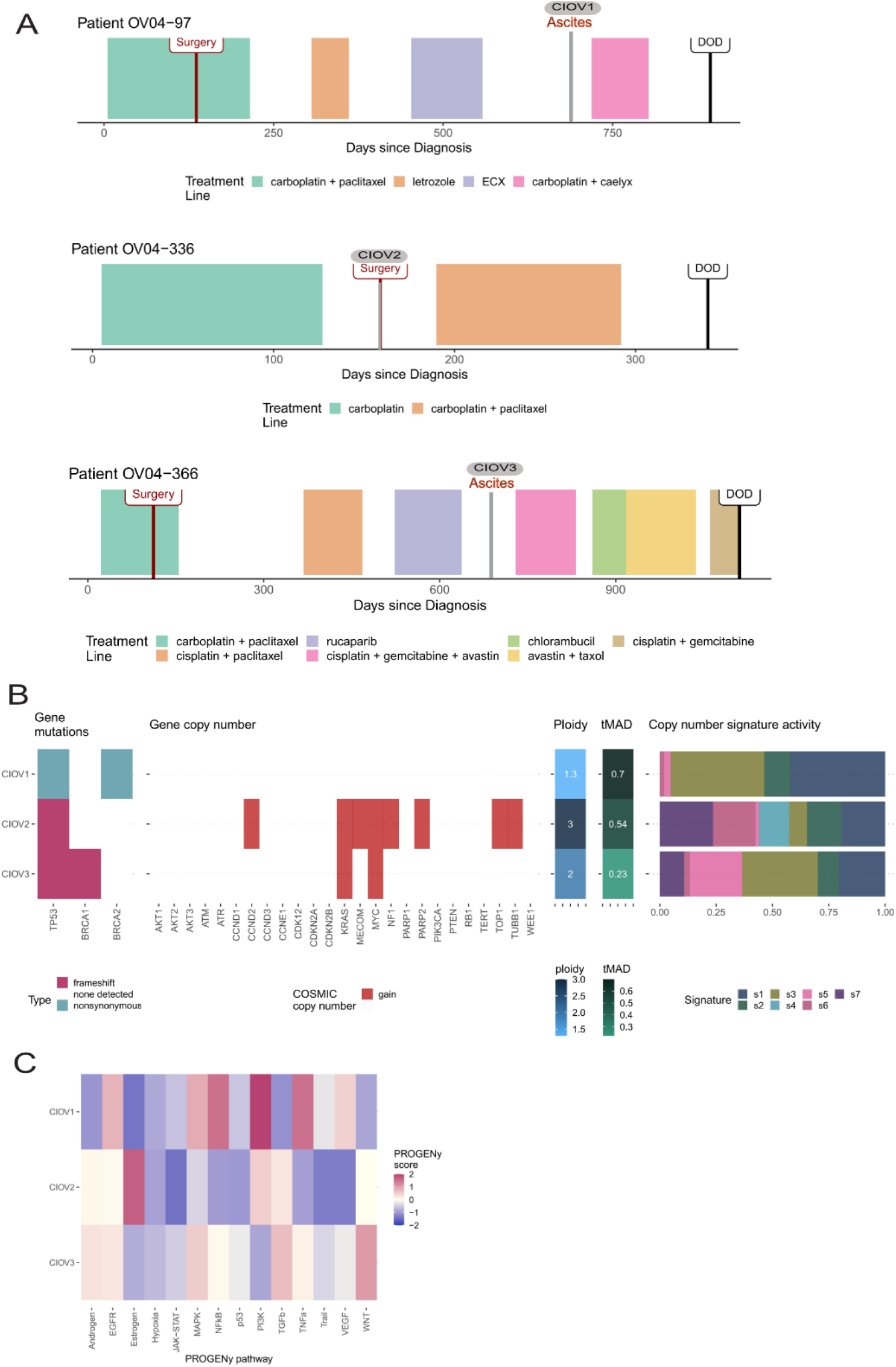
Clinical data and genomic characterization of cell lines. A. Chemotherapy regime for the three patients from which continuous cell lines were successfully generated. The time point where the ascites cells or the solid tumour were collected is represented by the name of the corresponding cell line in grey. B. Genomic characterization. Panel 1 Oncoprint plot of gene mutations called from TAm-Seq data. Panel 2 shows gene copy number states (amplification, deletion, gain or loss) for HGSOC genes of interest. Amplifications and deletions were called using the definitions outlined by the Catalogue of COSMIC. Amplifications: ploidy ≤2.7 and CN ≥5; or ploidy >2.7 and CN ≥9. Deletion:ploidy ≤2.7 and CN = 0; or ploidy >2.7 and CN < (ploidy − 2.7). CN gain and losses are defined as a CN change of +1 or −1 from a sample’s ploidy, respectively. Ploidies are indicated in panel 3. Trimmed median absolute deviation from CN neutrality (tMAD) scores are shown in panel 4. Panel 5 shows the copy number signature activities derived from ACN-fitted sWGS data for each cell line C.Copy number signature activities for the tumours and their corresponding cell lines. D. Heatmap of PROGENy signalling pathway activities.

At initiation of culture (p0) the samples were heterogeneous with a proportion of cells resembling a fibroblast-like shape. Long-term culture selected for uniform, epithelial-like cuboidal shaped cells (Supplementary Figure 1B). These cell lines will be made available to the researchers through ECCAC.

#### In vivo growth

The *in vivo* growth potential of the three cell lines was determined by subcutaneous injection into NSG mice (n=2 mice for each cell line). Only CIOV1 and CIOV3 cell lines formed tumours.

#### Cell line genotyping

To establish the origin of the cells and to ensure that no contamination with other cells occurred during the culturing process, Short Tandem Repeat (STR) profiling was performed using gDNA from the cell lines and its matched donor tumour tissue. STR profiling of the CIOV2 and CIOV3 cell lines matched their tumour of origin 100% whereas the CIOV1 cell line lost alleles 14 and 16 in loci D13S317 and vWA respectively. The loss of these two alleles gives an identity of 87.5% which falls within the ATCC recommended threshold for it to be related to the original tumour cells (Table 1).

**Table 1:** STR Genotyping.

|  | Amel | CSF1PO | D13S317 | D16S539 | D21S11 | D5S818 | D7S820 | TH01 | TPOX | vWA |
| --- | --- | --- | --- | --- | --- | --- | --- | --- | --- | --- |
| patient OVO4-97 | X | 10,11 | 11,14 | 11 | 30,32.2 | 12 | 10,11 | 9.3 | 8,12 | 16,17 |
| CIOV1 | X | 10,11 | 11 | 11 | 30,32.2 | 12 | 10,11 | 9.3 | 8,12 | 17 |
| patient OVO4-336 | X | 9,12 | 13 | 9,11 | 28,32.2 | 12,13 | 11 | 6 | 11 | 18 |
| CIOV2 | X | 9,12 | 13 | 9,11 | 28,32.2 | 12,13 | 11 | 6 | 11 | 18 |
| patient OVO4-366 | X | 10 | 12 | 12 | 30,31 | 12 | 10,12 | 9.3 | 8,11 | 16 |
| CIOV3 | X | 10 | 12 | 12 | 30,31 | 12 | 10,12 | 9.3 | 8,11 | 16 |

#### Mutation analysis

The new continuous cell lines and the original tumours were processed for mutation analysis by targeted sequencing. *TP53* alterations were identical between cell lines and their matched tumours: CIOV1 and its matched tumour contained a non-synonymous mutation (p.G524A) in exon 5 of *TP53* (Figure 2B and Table 2). Both the CIOV3 and its matched tumour contained a frameshift deletion (p.783delT) in exon 8 of *TP53* and CIOV2 and its matched tumour contained a frameshift deletion (p.A86Vfs*55) in exon 4 of *TP53*.

**Table 2:** Targeted sequencing results for tumours.

| Subject | Mutation Type | Gene | cDNA effect | Protein effect | Chromosome | Genomic location |
| --- | --- | --- | --- | --- | --- | --- |
| OV04-097 | non-synonymous | TP53 | c.G524A | R175H | 17 | 7578406 |
| OV04-336 | frameshift deletion | TP53 | c.257_279del23 | p.A86Vfs*55 | 17 | 7579407 |
| OV04-366 | frameshift deletion | TP53 | c.783del1 | S261Rfs*84 | 17 | 7577155 |
| OV04-366 | frameshift deletion | BRCA1 | c.4066del4 | N1355fs | 17 | 41243479 |
| OV04-366 | non-synonymous | BRCA1 | c.G4083A | M1361I | 17 | 41243465 |
| OV04-366 | nonsynonymous,splice_region | BRCA1 | c.G4096T | G1366C | 17 | 41243452 |

**Table 3:** Targeted sequencing results for CIOV cell lines at passage 20.

|  | Mutation Type | Gene | cDNA effect | Protein effect | Chrom | Genomic | MAF(%) |
| --- | --- | --- | --- | --- | --- | --- | --- |
| CIOV1 | nonsynonymous | TP53 | c.G524A | R175H | 17 | 7578406 | 99 |
|  | nonsynonymous | BRCA2 | c.8182G>A | p.V2728I | 13 | 32937521 | 99 |
| CIOV2 | frameshift deletion | TP53 | c.257_279del23 | p.A86Vfs*55 | 17 | 7579407 | 98 |
| CIOV3 | frameshift deletion | TP53 | c.783del1 | S261Rfs*84 | 17 | 7577155 | 98 |
|  | frameshift deletion | BRCA1 | c.4066del4 | N1355fs | 17 | 41243479 | 99 |
|  | nonsynonymous,splice_region | BRCA1 | c.G4096T | G1366C | 17 | 41243452 | 42 |

When cell lines were established (passage 20) we determined if they harboured mutations in a set of genes that are important in HGSOC: *PTEN, BRCA1, BRCA2, RAD51C, RAD51B, RAD51D, PALB2, NF1, CDK12, CTNNB1, FANCM, RB1, BRIP1, BARD1, EGFR, KRAS, NRAS*, and *PIK3CA*. Mutations above the cut-off threshold of 5% mutant allelic fraction included: a benign nonsynonymous mutation in *BRCA2* (Figure 2B) in CIOV1 and both a frameshift deletion p.4065-4068del4bp and a non-synonymous mutation p.G1366C in *BRCA1* in CIOV3 cell line. No mutations in this set of genes were detected in CIOV2.

#### Copy number signature activity

Analysis of the copy number alterations with low-coverage whole genome sequencing revealed a high degree of chromosomal instability in each cell line with patterns very similar to their parental patient tumour samples (Figure 2C and Supplementary Fig 2). Detailed genomics analysis of the cell lines has been already described (Sauer et al., 2023).

#### RNAseq and progeny analysis

We used the RNAseq data that we recently published (Sauer et al., 2023) to estimate PROGENy pathway activities in the three cell lines and found a marked increase in PI3K activity in CIOV1, estrogen pathways in CIOV2 and WNT signalling in CIOV3 (Figure 2C).

#### Immunostaining for p53 and EpCam

First, a pathologist verified H&E samples confirming the HGSOC histotype and similarity between tumours, cell lines and xenografts (Supplementary Figure 3A). Then, p53 expression in the solid tumours from patients OVO4-97, OVO4-366 and OVO4-336, cell lines and cell line-derived xenografts was assessed by immunohistochemistry and found that patient OVO4-97 was highly positive, patient OVO4-336 was negative and patient OVO4-366 showed some degree of cytoplasmic staining (Supplementary Figure 3B). Staining patterns in the cell lines and xenografts mirrored those of their parental tumour staining (Supplementary Figure 3B and 3C). To verify the epithelial origin of the cell lines we stained for EpCam expression levels and to identify mesenchymal cells we also stained for vimentin expression levels (Supplementary Figure 3D). Cell co-expressing Epcam and vimentin are likely undergoing epithelial to mesenchymal transition. CIOV1 was strongly positive for EpCam and negative for vimentin whereas CIOV2 and CIOV3 co-expressed EpCam and Vimentin.

#### Cell growth and oncogenic assays

The growth of the cell lines was determined using a live cell imaging system that takes images every 3 h and uses an algorithm to derive a measure of cell confluency. Doubling time of the cell lines was calculated from the proliferation curves: 25 h for CIOV1, 42 h for CIOV3 and 64 h for CIOV2. Cell proliferation was measured in both high (10%) and low (1%) serum (Supplementary Figure 4A). CIOV1 grew the fastest in both high and low serum, whilst CIOV2 grew more slowly and only in high serum. The migration potential of the cell lines was determined using a scratch wound healing assay, and we observed that CIOV1 cells migrated faster than CIOV3, whilst CIOV2 cells remained relatively static (Supplementary Figure 4B).

The capacity of the cell lines to form spheroids in 3D cultures was assessed as a surrogate of metastatic potential. Cells were grown in two different ways: in suspension using round ultra-low attachment plates (ULA) and embedded in Matrigel® growth factor reduced basement membrane matrix (ECM) and imaged 14 days after seeding (Supplementary Figure 4C). When grown in ULA plates CIOV1 cells formed loose cell aggregates whereas CIOV2 and CIOV3 formed compact spheroids. When grown embedded in ECM the three cell lines formed compact spheroids.

#### In vitro chemosensitivity

To determine the sensitivity of the cell lines to paclitaxel and cisplatin we measured the drug-induced growth rate (GR) inhibition. The GR value is defined as the ratio between growth rates under treated and untreated conditions normalized to a single cell division (Supplementary Figure 4D). At all concentrations cisplatin and paclitaxel were both cytotoxic for the CIOV1 cell line. Only at very high concentrations was cisplatin cytotoxic for CIOV2 and CIOV3, and partially cytostatic at lower concentrations. Paclitaxel is partially cytostatic at all concentrations tested therefore treated cells are growing but a slower pace than the untreated cells.

### Selection of cancer cells *in vitro*

To monitor clonal evolution during culture, we processed an aliquot of the CIOV1 and CIOV3 cell lines at some of the passages for genetic characterization. In a prior study (Schwarz et al., 2015), we demonstrated that patient OV04-97 had a subclonal population marked by an *NF1* deletion. However, in the derived CIOV1 cell line although present at passage 0, the *NF1* clone could not be detected at passage 3 (Figure 3A). By contrast, the *TP53* mutant allele fraction per cell increased concomitantly with increased passage number.

**Figure 3.**
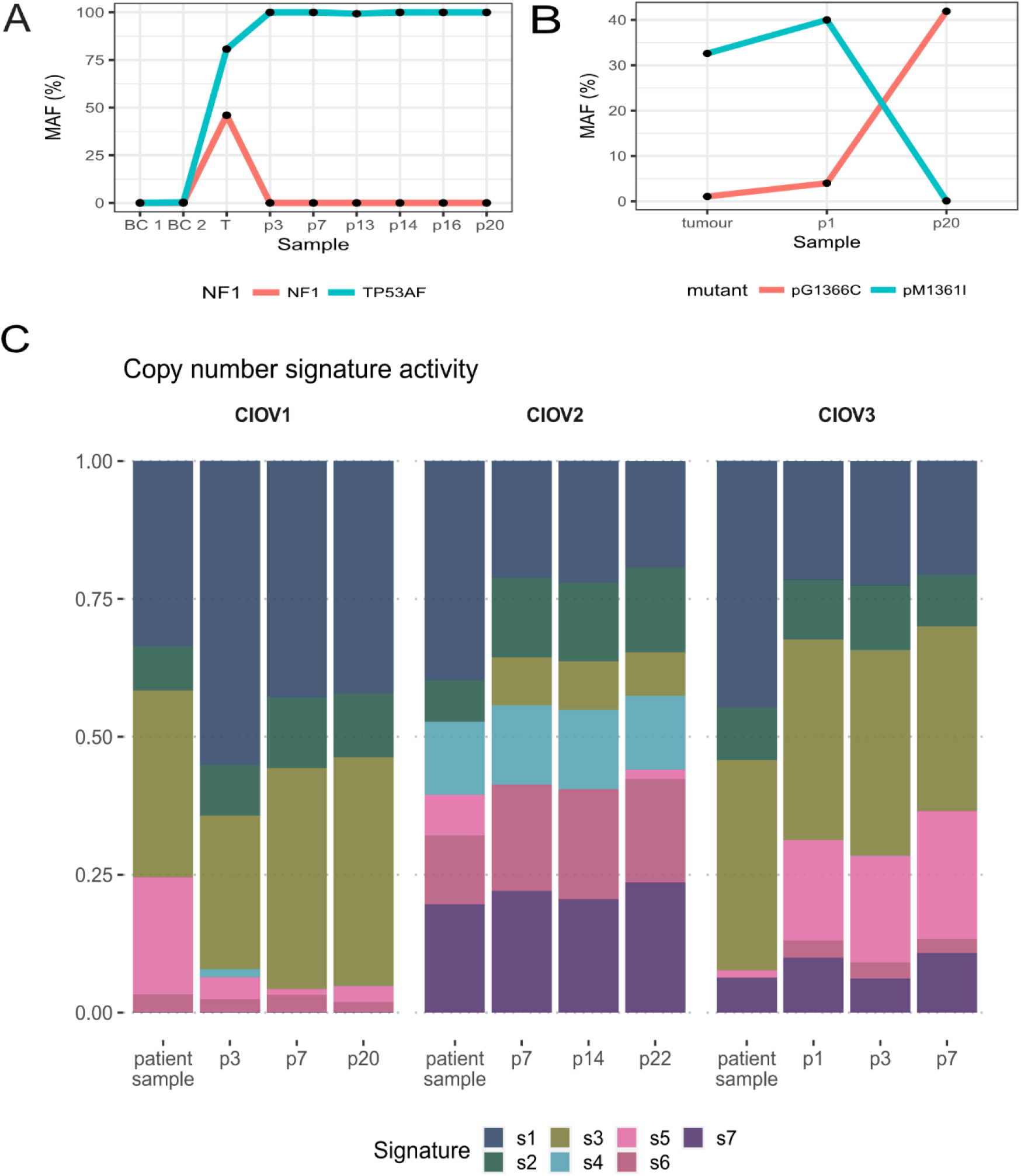
Cell selection in culture. A. *TP53* and *NF1* mutant allele fraction in tumour and CIOV1 cell line over time. B. *BRCA1* mutant allele fraction in tumour and CIOV3 cell line over time. C Copy number signatures for the tissue and the three cell lines at different passage numbers.

When monitoring clonal evolution of the CIOV3 cell line derived from patient OV04-366 we confirmed the presence of three BRCA1 mutations: p.N1355fs, p.G1366C and p.M1361I before culture. Variant calling analysis for each mutation revealed that the allele fraction for the p.G1366C mutation was very low in both tumour and cell line at passage 1 and increased to ∼42% at passage 20 (Figure 3B). The other mutation p.M1361I is high in the tumour and passage 1 but is found at 0.1% in passage 20. Although we observed some selection at the mutational level, the analysis of cell line copy-number signatures over time showed (Figure 3C) no significant changes between passages.

## Discussion

The heterogeneity of high grade serous ovarian cancer is a significant clinical challenge that must be overcome to extend patient survival. Reliable *in vitro* model systems representing the full spectrum of genetic modifications observed in patients are therefore essential for our research. This caveat was already described by Domcke et al. (Domcke et al., 2013) for many of the widely used ovarian cancer continuous cell lines where clinical data was poor. Other papers describing the establishment of HGSOC cell lines have been published (Fleury et al., 2015; Kreuzinger et al., 2015; Thu et al., 2017) but although they used the same culture conditions for all the cells there is no comprehensive evaluation of the time or number of tissue samples needed to generate the cell lines. Another major challenge to make continuous cell lines is to find adequate media formulation for each genetic background; to address this Ince et al made complex media and reported 95% success rates (Ince et al., 2015) but culture conditions were not standardised in terms of plasticware and oxygen levels which made comparisons difficult. In this paper, we have systematically grown 114 primary cell lines from solid tumours, pleural fluid, and ascites that we obtained from our consented patients and demonstrated that under the same culture conditions not only 35% of the tumour cells die at passage 0 but also most cells senesce before becoming a continuous cell line. HGSOC finite cell lines can be successfully generated but with the consequent limitation of having a short lifespan making them unsuitable for certain experiments and compromising reproducibility across laboratories. The low rates achieved in generating continuous HGSOC cell lines might explain the low number of cell lines of this ovarian cancer subtype available to-date as compared to other tissue types.

In this study, three spontaneously immortalized continuous cell lines have been generated and fully characterized. During the establishment process cells growing in 2D culture conditions initially adapt and our data suggests that some changes occurred at early passages probably due to cell heterogeneity but overall, they resemble the tissue of origin. The epithelial origin of the cell lines was confirmed by EpCam expression, but interestingly CIOV2 and CIOV3 also expressed vimentin suggesting an epithelial-mesenchymal transition in these cell lines. To account for the differences seen in the growth rate and doubling time of the cell lines we applied the GR metrics to the drug sensitivity test. Although speculative, it is intriguing that the cell lines that present an EMT phenotype (CIOV2 and CIOV3) are resistant to conventional chemotherapy agents, consistent with studies showing that EMT is associated with drug resistance (Zhang and Weinberg, 2018; Dongre and Weinberg, 2019; Yang et al., 2020).

Although we have generated and fully characterised three continuous cell lines the success rate was poor and the method laborious. Further optimization and new technologies are needed to generate the cell lines required to represent the heterogeneity of HGSOC.

## Conclusion

We have shown that growing high grade serous ovarian cancer cells as 2D-continuous cell lines using standard media is a very challenging process. Although the establishment rate was low, we generated three new continuous high grade serous ovarian cancer cell lines. Using TAM-seq, IHC, and shallow WGS we confirmed that each cell line retains the genomic and pathologic characteristics of the parental tumour. These new HGSOC tumour models will become useful tools for the research community studying chemotherapy resistance and sensitivity.

## Author contributions

Conceptualization, M.V. and J.D.B.; Methodology, M.V.; Investigation, M.V., C.M.S., I.G., and C.C.; Resources, M.V., C.M.S., I.G., T.G, E.V.O., D.H. and A.P.; Curation of clinical data, E.M and G.F.; Writing-Original Draft, M.V.; Writing-Review and Editing, M.V., C.M.S., H.M.E. and J.D.B.; Visualization, M.V. and C.M.S.; Supervision, M.V. and J.D.B.; Project Administration, M.V.; Funding Acquisition, J.D.B.

## Acknowledgements

We thank all patients who participated in and donated tissue samples to this study. We also thank Karen Hosking, Mercedes Jimenez-Linan and the OV04 study team for their help with clinical tissue samples. We would like to thank the Cancer Research UK Cambridge Institute Genomics, Biological Resource Unit, Compliance & Biobanking, Research Instrumentation and Cell Services, and Bioinformatics core facilities for their support with various aspects of this study. We would like to thank Nicholas Clark for his advice with the GRmetrics analysis.

