## supplementary figures for "Primary culture of high grade serous ovarian cancer cells, selection and derivation of three cell lines"

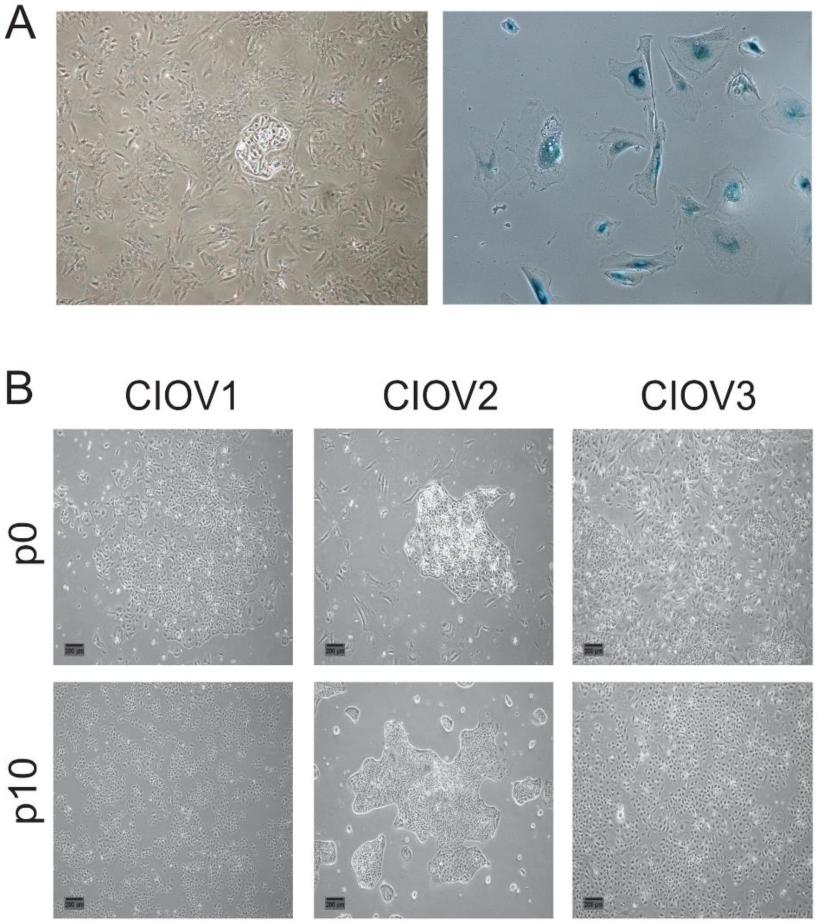


**Supplementary Figure 1. A. Cell culture complications**. Left panel: representative image of a tumour clone surrounded by fibroblast cells. Right panel: representative image of senescent cells. B. Cell morphology changes. Representative images of the cells in culture at passage 0 and 10. At p0 the samples were a mixture of epithelial and fibroblast-like cells whereas at later passages the cell population was morphologically homogenous showing a cobblestone-like morphology.

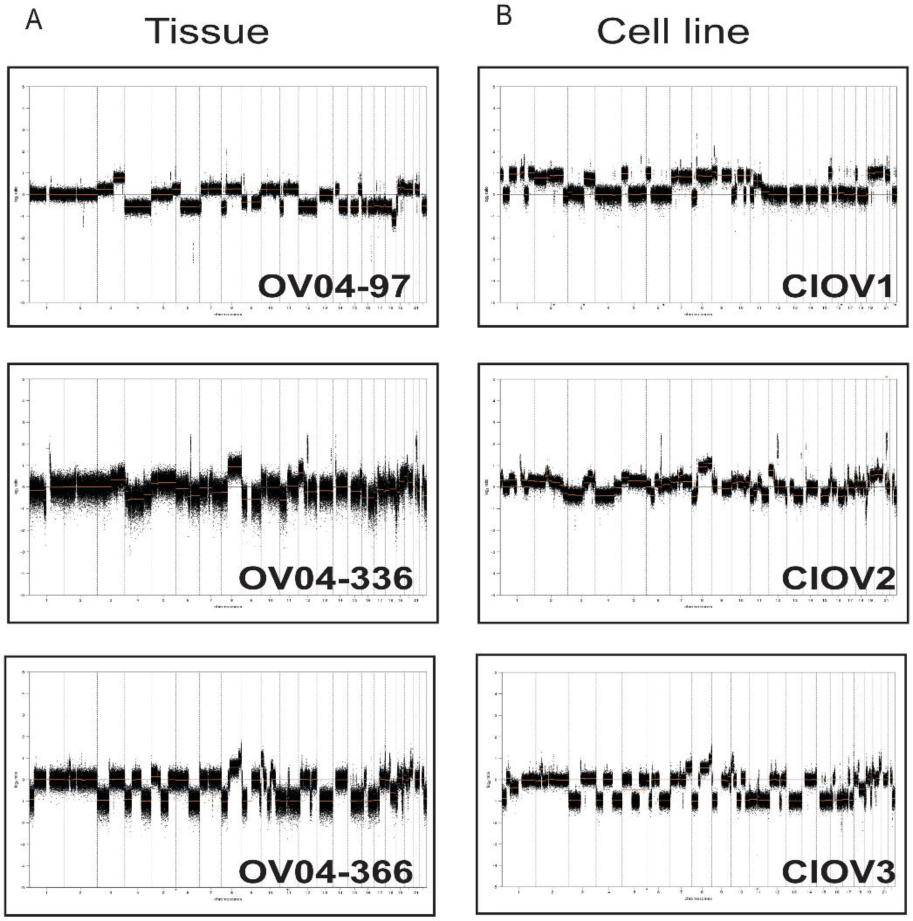


**Supplementary Figure 2. Gene copy number aberration plots**. A. Plots for Tissue samples B. Plots from established cell lines

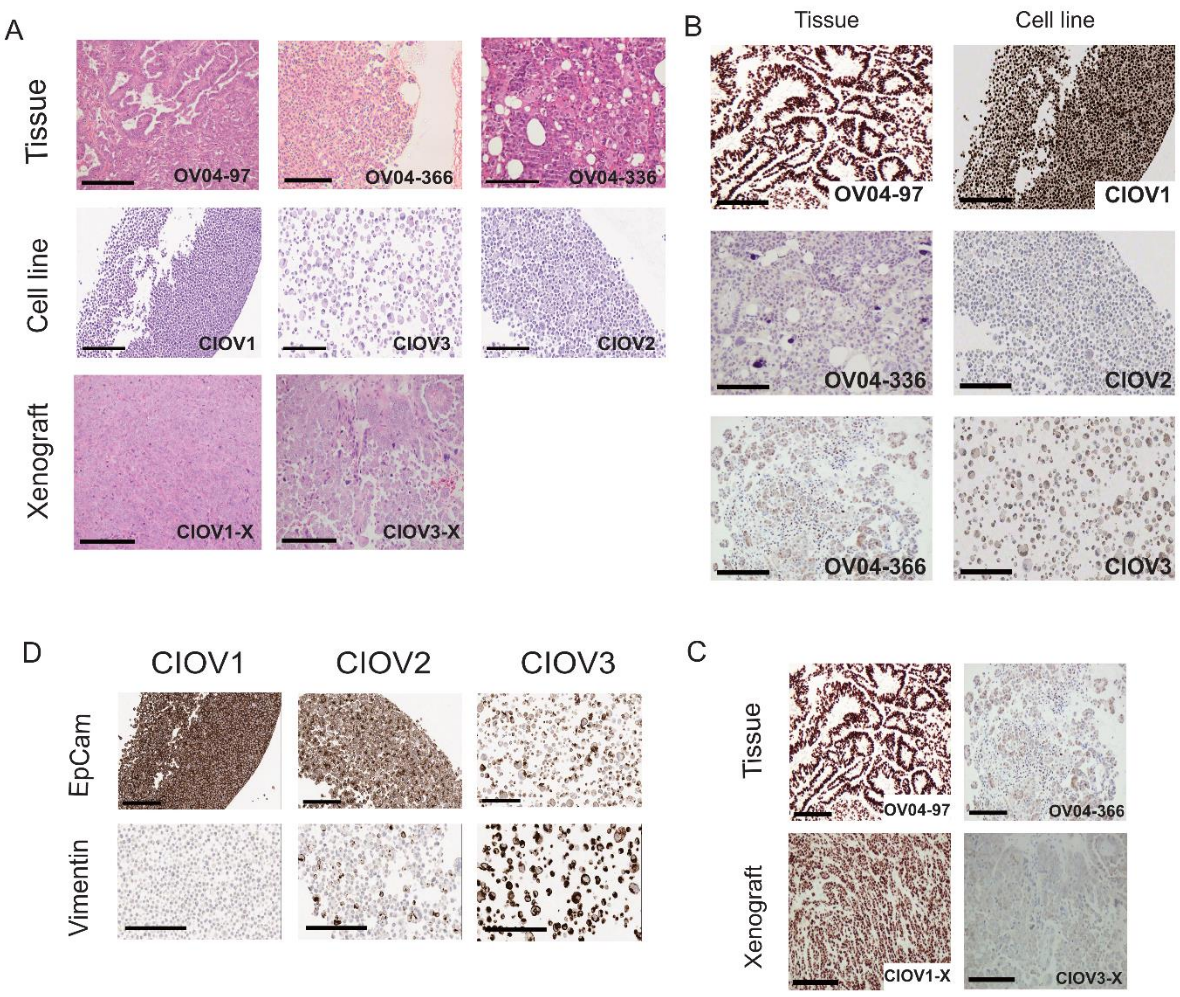


**Supplementary Figure 3** A. H&E staining of the tumours, cell lines and cell line-xenografts. Scale bars represent 200µm. B. Immunohistochemistry for p53 in the tumour tissue sections and the cell lines. Scale bars = 200µm. C. p53 immunostaining of the tissue and corresponding xenograft. D. EpCam and Vimentin immunostaining in the cell lines.

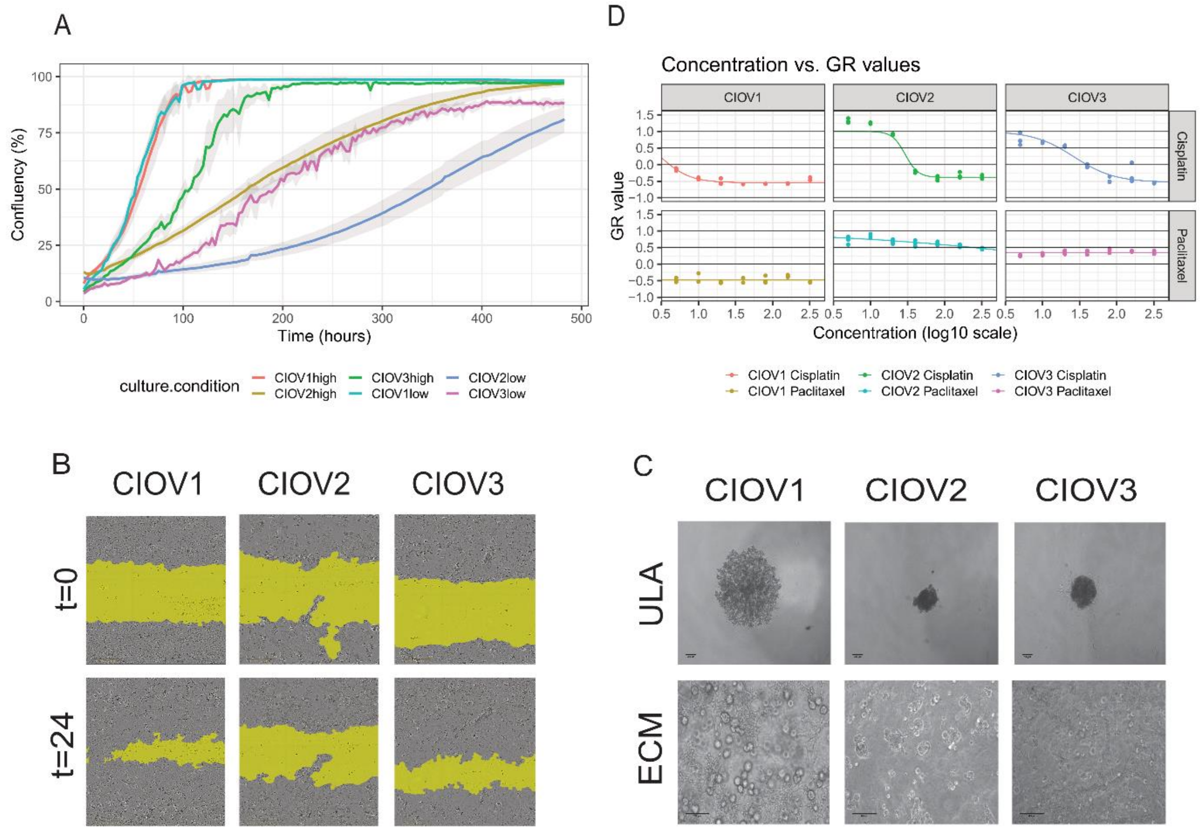


**Supplementary Figure 4**. A. Cell proliferation assay in both high and low serum content. B. Scratch wound healing assay. Relative wound density was calculated using the Incucyte metrics. C. Cells grown in 3D. Cell lines were grown in suspension in ultra-low attachment plates (ULA) or embedded in Matrigel (ECM). D. Chemosensitivity of the cell lines to paclitaxel and cisplatin. Assay was performed using GR metrics.
